# Lower cortical *FKBP5* DNA methylation at key enhancer sites is associated with older age and higher gene expression in schizophrenia

**DOI:** 10.1101/2025.01.28.635384

**Authors:** Katrina Z. Edmond, Dominic Kaul, Natan Yusupov, Maik Ködel, Susann Sauer, Anna S. Fröhlich, Ran Tao, Joel E. Kleinman, Daniel R. Weinberger, Thomas M. Hyde, Darina Czamara, Elisabeth B. Binder, Natalie Matosin

**Affiliations:** The University of Sydney, School of Medical Sciences, Sydney NSW 2050; The University of Sydney, Charles Perkins Centre, Sydney NSW 2050; Molecular Horizons, School of Chemistry and Molecular Bioscience, Faculty of Science, Medicine and Health, University of Wollongong, Northfields Ave, Wollongong 2522, Australia; Department of Genes and Environment, Max-Planck Institute of Psychiatry, Munich, Germany; International Max Planck Research School for Translational Psychiatry (IMPRS-TP), Munich, Germany; The Lieber Institute for Brain Development, Johns Hopkins University Medical Campus, Baltimore, MD, USA; Department of Psychiatry and Behavioral Sciences at Johns Hopkins University School of Medicine; Department of Psychiatry and Behavioral Sciences, Emory University School of Medicine, Atlanta, USA

**Keywords:** FKBP5, FKBP51, DNA methylatio, Epigenetics, Psychosis, Depression, Ageing, Stress, Single cell, Postmortem brain

## Abstract

An increasingly compelling body of literature indicates the glucocorticoid receptor cochaperone FK506-binding protein 51 (FKBP51) is a promising target for novel psychiatric therapeutics. However, the mechanisms regulating the corresponding *FKBP5* gene directly in the human brain remain largely unknown yet are needed to facilitate the development of precise mechanism-based treatment approaches. Here, we examined *FKBP5* DNA methylation patterns in postmortem human brain samples from the dorsolateral prefrontal cortex of individuals who lived with a major psychiatric disorder (schizophrenia, major depression, or bipolar disorder; n=329) and controls n=231. We identified that cytosine-phosphate-guanine-dinucleotide (CpG) specific *FKBP5* DNA methylation is altered in psychiatric disorders across the *FKBP5* locus, and that these changes are differentially associated with age and genotype (rs1360780 CC *vs* CT/TT). Individuals with schizophrenia had significantly lower levels of DNA methylation in the proximal enhancer of *FKBP5*, which also negatively correlated with *FKBP5* gene expression. These changes were also associated with predicted glucocorticoid response elements (GREs) in the proximal enhancer, but not other transcription factor binding sites. This evidence supports that in the human cortex, *FKBP5* DNA methylation is associated with both genetic and ageing effects, and that the associations between these factors vary at a diagnosis-specific level in psychopathology. This may have implications for developing *FKBP5*-targeted therapeutics and defining a subgroup of patients who will benefit from such treatments.

## INTRODUCTION

As the leading contributor to global years lived with disability [8], psychiatric disorders are a major global health burden. The approved use of available drugs for specific psychiatric diagnoses, as well as their patient-specific prescription, is currently based on attenuating primary symptoms rather than addressing the underlying biological mechanisms contributing to heterogenous disease presentation [11, 29, 43]. An improved understanding of the genetic and molecular underpinnings of psychiatric disorders can facilitate mechanism-based diagnoses of biologically distinct patient subgroups, including the corresponding biological targets for their treatment [43].

One mechanism-based target is the allosteric co-chaperone of the glucocorticoid receptor (GR) FK 506 Binding Protein 51 (FKBP51; *FKBP5* chromosome 6p21.31), a critical actor in propagating and terminating the stress response [52]. Human genetic, epigenetic and animal studies indicate that heightened *FKBP5* expression, particularly in individuals exposed to early life adversity, may contribute to increased psychiatric disease risk [30, 52]. Small-molecule FKBP51 antagonists show promising preclinical results, with mice administered FKBP51 antagonists, either systemically or directly into relevant brain regions such as the amygdala, having improved stress-coping behaviour and reduced anxiety [11, 15]. To refine and further develop this pharmacological approach, understanding the cell- and region-specific pathological changes to *FKBP5* transcription and drivers behind its regulation is crucial.

*FKBP5* induction is shaped by an interplay of clinically relevant genetic and epigenetic mechanisms [30]. A minor allele of a functional *FKBP5* common haplotype (single nucleotide polymorphism [SNP] rs1360780 T allele in intron 2) is associated with exacerbated *FKBP5* induction following stress or glucocorticoid exposure [22]. Carriers of the *FKBP5* risk haplotype exposed to childhood adversity are at an increased risk for developing psychopathology [30]. Reduced DNA methylation (DNAm) at key *FKBP5* glucocorticoid response elements (GREs) is reported following exposure to glucocorticoids [48]; and this was associated with higher *FKBP5* mRNA induction in both human peripheral blood samples and a multipotent human hippocampal progenitor cell line [22, 23]. These two mechanisms are hypothesised to converge in a subgroup of individuals who carry the *FKBP5* minor allele haplotype and were exposed to early life adversity [22, 23]. An increase in *FKBP5* induction following stress exposure in risk allele carriers leads to a prolonged glucocorticoid release via direct effects of *FKBP5* on GR function and stress hormone axis regulation [22, 23]. Genetic predisposition combined with exposure to early life adversity may thus contribute to a joint genetic and epigenetic disinhibition of *FKBP5* transcription in response to stress [30], raising risk to psychopathology [20].

*FKBP5* expression levels also consistently increase over the life course, even in healthy people [3, 12, 30, 31, 47]. This developmental pattern of expression may be modulated by *FKBP5* DNAm in regulatory regions of the gene, including GREs in intron 2 and 7 [3, 37]. Environmental exposures such as early life adversity also impact on DNAm to heighten the *FKBP5* ageing trajectory in the risk haplotype carriers [30]. This convergence of risk factors – ageing and early life adversity on heightened *FKBP5* responsivity – has been reported in peripheral blood immune cells, with increased FKBP51 contributing to a proinflammatory profile with increased age via activation of the alternative nuclear-factor _K_B pathway [53]. The next step is understanding if and how this translates to the brain, where psychiatric symptoms manifest, and targeted therapeutics will exert their primary effects.

In this study, we comprehensively investigated associations of *FKBP5* DNAm on *FKBP5* expression in the neocortex [31]. We used DNAm arrays, SNP genotyping and bulk RNA sequencing on a large collection of postmortem human dorsolateral prefrontal cortex samples (n=560, DLPFC; BA9) from individuals that lived with schizophrenia, major depression and bipolar disorder and matched controls. We demonstrate age- and genotype-mediated diagnosis-specific alterations in *FKBP5* DNAm and replicate these findings in an independent cohort (orbitofrontal cortex [OFC], BA11, n=91). Interestingly, these effects converge most significantly in cytosine-phosphate-guanine-dinucleotides (CpGs) located within GREs of the proximal enhancer region and the topologically associated CCCTC-binding factor (CTCF) sites, especially in older individuals with schizophrenia. Thus, CTCF associated CpGs within the proximal enhancer of *FKBP5* are genomic regulatory regions likely to directly impact on gene expression levels.

## MATERIALS AND METHODS

### Human postmortem brain samples

*FKBP5* gene expression, DNAm and SNP genotype were examined in the DLPFC (BA9) of 560 postmortem human brains acquired by the Lieber Institute for Brain Development Repository via material transfer agreement (Table 1, cohort 1). Brain tissue was donated at the time of autopsy from legal next of kin using audiotaped witnessed informed consent, through the NIMH Human Brain Collection Core under NIMH/NIH Institutional Review Board protocol #90-M-0142, and through the National Institute of Child Health and Human Development Brain and Tissue Bank for Developmental Disorders (http://www.BTBank.org/), under contracts NO1-HD-4-3368 and NO1-HD-4-3383, approved by the Institutional Review Board of the University of Maryland. Subjects included a lifetime cohort of non-psychiatric controls (n=340), spanning in age from the prenatal second trimester (14weeks) to 85 years, in addition to teenagers, adults, and 50+ year old subjects with schizophrenia (n=121), major depression (n=144) or bipolar disorder (n=63). Detailed cohort demographics and information regarding tissue collection and dissection procedures have been published previously [31]. DNAm and SNP genotype of the OFC (BA11) were available to examine in a replication cohort (Table 1, cohort 2) of 91 subjects from the NSW Brain Tissue Resource Centre (University of Sydney, Australia). The study was approved by the Ludwig Maximilian University Ethics Committee (project 17-085, application 22-0523) and prospective informed consent by the donor, or legal next of kin was obtained for each subject. Detailed cohort demographics and information regarding tissue collection and dissection procedures have been published previously [31].

**Table 1:**
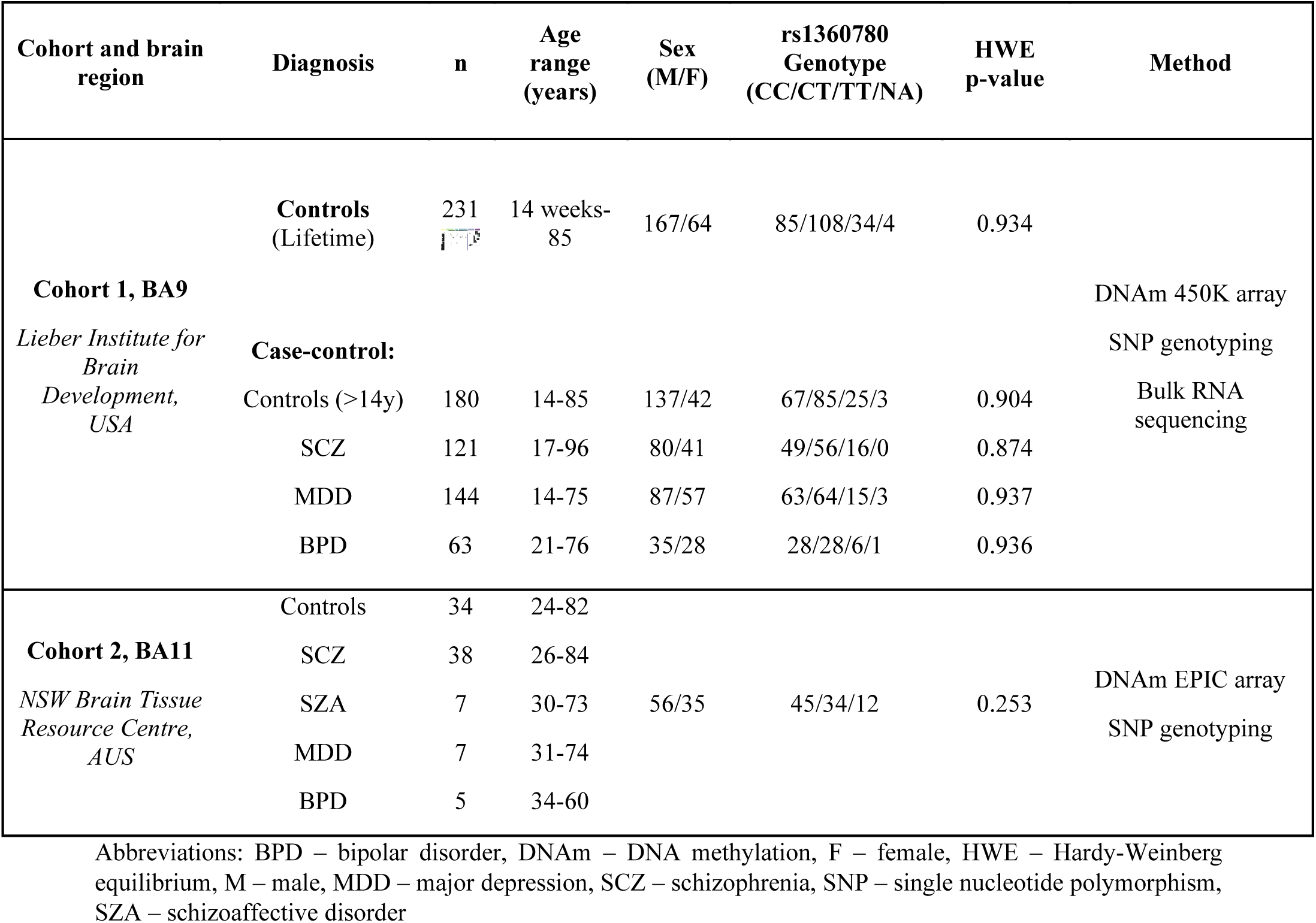
Demographics of cohorts used in study.

### DNA methylation microarray

Methods for analyses of DNAm have been previously described [10, 17]. Briefly, for cohort 1, genomic DNA for each subject was extracted from 100mg of pulverized DLPFC tissue using phenol-chloroform. 600ng of genomic DNA was bisulfite converted with the EZ-DNA methylation kit (Zymo Research, Irvine, CA, US). DNAm was assessed using the HumanMethylation450 BeadChip kit (Illumina, San Diego, CA, USA) which covers >485,000 CpG dinucleotides covering 99% RefSeq gene promoters, including the *FKBP5* gene promotor. Arrays were run according to the manufacturer’s protocols, with a percentage of the samples run in duplicate across multiple processing plates for assessing technical variability due to DNA extraction and bisulfite conversion. DNAm data was processed and normalized using Bioconductor’s Minfi package in R [1]. Batch effects were corrected as previously described, with batch-derived principal components used as covariates [17]. In cohort 2, genomic DNA for each subject was extracted from 10mg of frozen OFC tissue using QIAamp DNA mini kit (Qiagen, Hilden, Germany) following the manufacturer’s instructions, and concentrated using DNA Clean & Concentrator-5 (Zymo Research, Irvine, CA, US). 400ng of genomic DNA was bisulfite converted with the EZ-96 DNA Methylation kit (Zymo Research, Irvine, CA). DNAm was assessed using the Illumina Infinium MethylationEPIC BeadChip (Illumina, San Diego, CA, USA) according to manufacturer’s guidelines. DNAm data was processed and normalized (stratified quantile normalization and subsequent beta-mixture quantile normalization)[28, 41], using Bioconductor’s Minfi package in R [1]. Batch effects (array and row) were corrected with ComBat of the sva R package [25], while further brain tissue-related variables, which significantly correlated with the first five PCs (brain pH and storage time) were included as covariates in all analyses as previously described [10]. Cell-type composition was estimated for relative proportions of cell-types (neuronal [NeuN+] and non-neuronal) for both cohorts using epigenome-wide DNAm data as previously described [10, 17].

### SNP genotyping

For cohort 1, genomic DNA for each subject was extracted from 100mg of pulverized cerebellum tissue with the phenol-chloroform method. SNP genotyping was performed with the HumaHap650Y_V3 or Human 1M-Duo_V3 BeadChips (Illumina) according to manufacturer’s instruction and underwent quality control as previously described [39]. For cohort 2, genotyping was conducted using Illumina global screening arrays (GSA-24v3-0, Illumina, San Diego, CA, US) according to manufacturer’s instruction and underwent quality control as previously described [10]. Allelic information for the rs1360780 SNP was extracted and Hardy-Weinberg equilibrium (HWE) p-values calculated for both cohorts (Table 1).

### RNA sequencing

RNA sequencing methods have been previously described [39]. Briefly, total RNA for each subject was extracted from 100mg pulverized DLPFC tissue using RNeasy Lipid Tissue Mini Kits (Qiagen, Germantown, MD, USA). RNA was purified with RNeasy Mini Spin columns including on-column DNase digestion (Qiagen). Total RNA yield was determined with Qubit (ThermoFisher Scientific, Waltham, MA, USA) and RNA quality and RNA integrity assessed with the Agilent Bianalyzer 2100 (Agilent Technologies, Santa Clara CA USA). Poly-A containing RNA was purified from 1μg total RNA, and mRNA molecules fragmented using divalent cations and heating. cDNA conversion was achieved with reverse transcriptase and random hexamers, and second-strand cDNA was synthesized with DNA polymerase 1 and RNase H. T4 DNA polymerase, T4 polynucleotide kinase (PNK0), and Klenow DNA polymerase were applied for end-repair. The addition of an ‘A’ base was achieved using Klenow Exo (3’ to 5’ exon minus) and ligation of Illumina P adapters to allow for pooling <8 samples in one flow cell. Purification and enrichment were then achieved by polymerase chain reaction (PCR) to create the final cDNA library. High-throughput sequencing was performed on the HiSeq 2000 (Illumina), with the Illumina Real Time Analysis module used for image analysis and base-calling, and the BCL Converter (CASAVA v1.8.2) to generate FASTQ files with sequencing pair-end 100 base pair (bp) reads.

Splice-read mapper TopHat (v2.0.4) was used to align reads to the human genome reference (UCSC hg19), with known transcripts provided by Ensembl Build GRCh37.67. Mapped reads covering the genomic region of *FKBP5* (chr6:35541362-35696397, GRCh37/hg19) were acquired. Reads covering each exon or unique exon-exon junction level were called using featureCounts (V1.5.0) [26]. Individual raw exon and junction reads were divided by the mapped total reads per subject. To remove residual confounding by RNA degradation, we used a quality surrogate variable analysis (qSVA) framework as previously described [18].

### Transcription factor binding regions

To assess overlap of transcription factor binding sites with *FKBP5* CpGs we considered non redundant peaks for hg19 (lifted) stored in the ReMap 2022 catalogue, a large-scale integrative catalogue of DNA-binding experiments (https://remap.univ-amu.fr/)[14]. Catalogue was subset for exact overlap with genomic coordinates of investigated *FKBP5* probes (n=50) on the DNAm array using the subsetByOverlaps function of the IRanges R package [24]. Results from experiments related to the human cerebral cortex were further evaluated. Binding sites were derived from two human cortex datasets GSE129039 [46] and GSE116825 [27].

### Statistical methods for case-control and ageing analyses

All statistical analyses were performed in R v4.4.1 (cohort 1) and v4.0.4 (cohort 2) (https://www.r-project.org) [35]. For main effects (cohort-wide) on DNAm, a linear regression model was fit across the whole of cohorts 1 and 2 (cases and controls). For cohort 1 age, sex, genotype (rs1360780 CC vs CT/TT), brain pH, postmortem interval (PMI), estimated proportion of neuronal cells (NeuN+), the first four principal components estimated from the background probes (negative control), and the principal components of genotype that affected the main model were included to account for ancestry. For cohort 2, age, sex, genotype (rs1360780 CC vs CT/TT), brain pH, storage time, estimated proportion of neuronal cells and three first principal components of genotyping data (ancestry) were included. Cohort 2 was not used for case-control analysis across the *FKBP5* locus given the significantly smaller sample size. Significant CpGs for each covariate were assessed (P_FDR_ <0.05). For case-control analysis, a linear regression model was fit for all tested CpGs and gene expression correcting for age, sex, genotype (rs1360780 CC vs CT/TT), brain pH, PMI, estimated proportion of neuronal cells (NeuN+), the first four principal components estimated from the background probes (negative control), and the principal components of genotype (ancestry) that affected the main model. For case-control or genotype by ageing analysis, the R package *sm* was used to test for equality of non-parametric regression curves available in the *sm.ancova* command [4, 50]. As *sm.ancova* does not accept covariates, we calculated the residuals of a linear model accounting for covariates with these residuals then used this as response values in *sm.ancova*. All case-control analyses were restricted to subjects 14-85 years given the youngest case (major depression) was 14 years old and the oldest case was 81 years old. Multiple-testing correction was applied using a false discovery rate (FDR) cut-off value of 0.05 [2], unless otherwise noted.

## RESULTS

### Age and genotype effects on cortical *FKBP5* DNAm

Prefrontal cortex (PFC)-mediated signalling is an essential component of cognitive control and executive functioning [32]. Given the association of PFC regions, such as the DLPFC and OFC, with psychiatric phenotypes [13, 19] and altered trans-diagnostic *FKBP5* expression [31, 38, 40], we focused on profiling cortical *FKBP5* DNAm changes. To investigate the functional significance of changed DNAm across *FKBP5*, we first annotated boundaries functionally associated with the *FKBP5* gene, specifically the distal and proximal topologically associating domains (TADs) (∼155kb coverage, chromosome 6p21.31; Figure 1a-i). We then assessed all CpGs covered by the Illumina HumanMethylation450 BeadChip (450K) within these TAD boundaries, detecting a total of 50 CpGs (Figure 1a-ii). These CpGs were annotated based on functional domains, including the distal TAD, a downstream conserved DNAm quantitative trait locus, intron 5, the transcription start site, the proximal enhancer, and the proximal TAD (Figure 1a, Supplementary Table 1).

**Figure 1:**
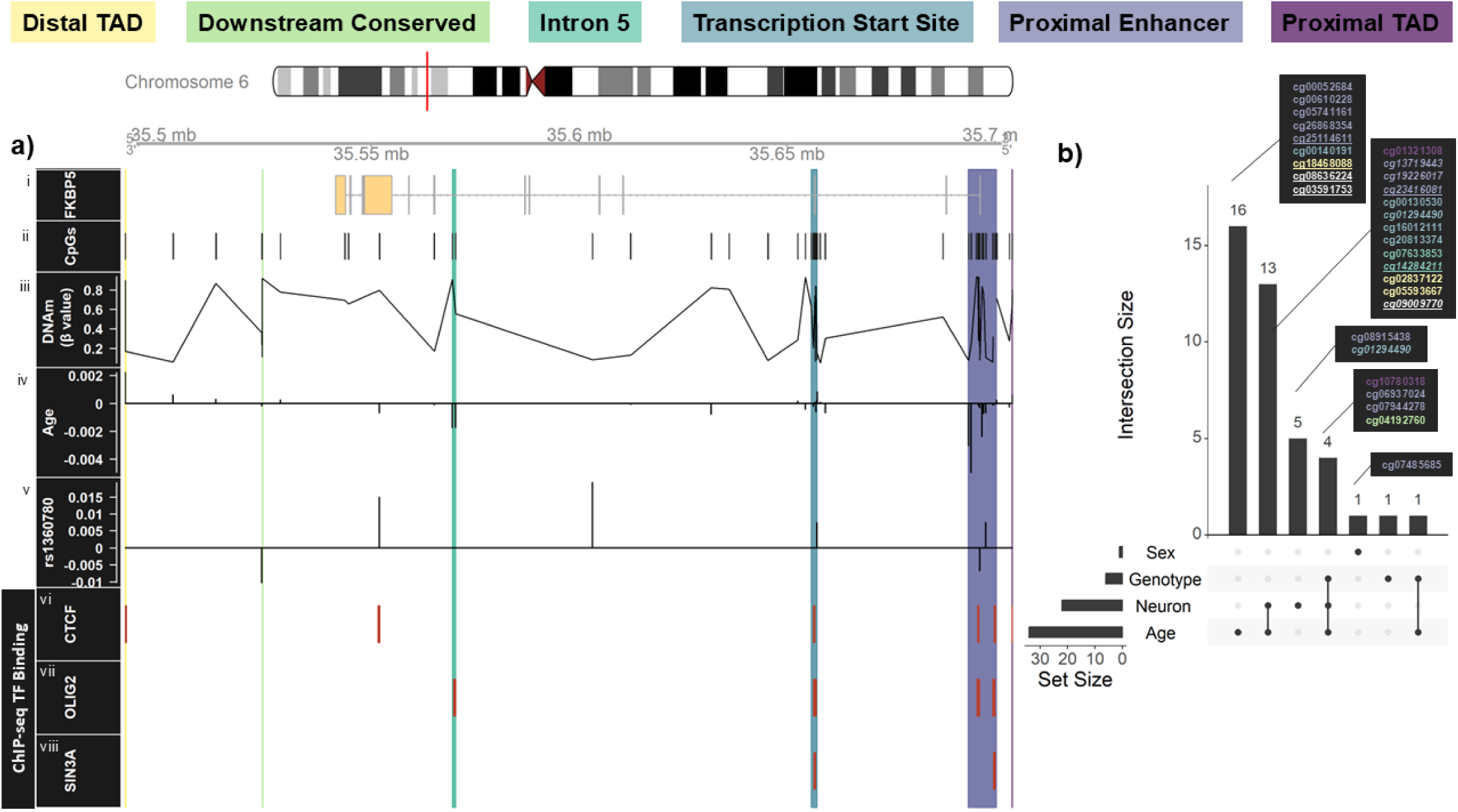
Functionally annotated CpGs in *FKBP5* locus demonstrate strong ageing and genotype effects across lifespan. **(a)** Annotation of the *FKBP5* locus. Highlighted domains indicate functionally annotated domains. (i) *FKBP5* gene map located on chromosome 6. (ii) All CpGs captured using the 450K Array. (iii) Mean beta (β) value (DNAm) across cohort. (iv) CpGs significantly associated with age (P_FDR_<0.05). Line direction relative to baseline indicates mean effect size. (v) CpGs significantly associated with genotype (rs1360780) (P_FDR_<0.05). Line direction relative to baseline indicates mean effect size. (vi-viii) ChIP-seq binding sites of transcription factors using the ENCODE database (https://www.encodeproject.org/), restricted to studies of the human cortex (GSE129039 [46] and GSE116825 [27]). **(b)** UpSet plot representing intersection of significant CpGs. Sets connected by a line indicate overlapping CpGs and the height of the column indicates the size of the overlap. Individual dots are unique CpGs to a given set. Callouts listing CpGs are coloured by functionally annotated domain. CpGs indicated with an <u>underline</u> were ageing effects conserved in the replication cohort (BA11) P_FDR_<0.05. Abbreviations: CpG – cytosine-phosphate-guanine-dinucleotides, TAD – topologically-associated domain, TF – transcription factors.

Previous studies consistently show increased expression of *FKBP5* across age and in rs1360780 genotype minor allele (T) carriers [30].

**Age:** In cohort 1 (DLPFC, n=329 cases and n=208 controls, 0-96 years), we identified age to be broadly associated with altered DNAm in *FKBP5* (Figure 1b), with 34 (68%) of the analysed CpGs showing significantly altered levels of DNAm (FDR corrected; Figure 1a-iv, 1b). Over half of these CpGs (n=21) fell within the six functionally annotated regions of *FKBP5* and ageing was generally associated with reduced DNAm. The proximal enhancer was the primary contributor with 10 significant CpGs (cg23416081, cg00052684, cg00610228, cg17030679, cg25114611, cg19226017, cg08915438, cg05741161 cg26868354, cg13719443; Figure 1a-iv). The remaining 11 significantly altered CpGs were split across four of the five remaining functional regions, specifically, three in the distal TAD (cg05593667, cg02837122, cg18468088), two in intron 5 (cg07633853, cg14284211), five in the transcription start site (cg00140191, cg16012111, cg01294490, cg20813374, cg00130530), one in the proximal TAD (cg01321308) and none in the downstream conserved region (Figure 1a-iv). We also confirmed that these trends persisted in functionally annotated regions in an extended dataset (n=180 + 51) of cohort 1 healthy controls spanning prenatal to 98 years age, with increased DNAm in the distal TAD and decreased DNAm in the proximal enhancer observed across ageing (Supplementary Figure 1). To validate these results, we replicated our analysis using an EPIC array dataset derived from the OFC of healthy controls and subjects with psychiatric disorders (cohort 2, Table 1). 41 out of the 50 CpGs from cohort 1 were captured on the EPIC array, cg08915438, cg03566752, cg18726036 were not included and the remaining six did not pass quality control. Five of the six excluded CpGs were significant for ageing effects in cohort 1. Of those which were included, effects for four CpGs associated with ageing in cohort 2 were consistent with cohort 1 (Figure 1b). As in cohort 1, the proximal enhancer was the primary site of altered DNAm (cg23416081, cg25114611). Spearman’s correlation identified a strong positive association between the directionality of ageing effects between the two cohorts (rho=0.56, p=0.004).

**Genotype:** Given that genotype alters the 3D structure and indirectly affects DNAm in *FKBP5* [52], we also explored the impact of genotype. Our analyses identified a significant association of the rs1360780 genotype with DNAm levels in *FKBP5* (Figure 1a-v, Figure 1b). Of the 50 CpGs analysed, six showed significantly altered levels (FDR corrected) of DNAm in CT/TT allele carriers compared to non-risk haplotype CC carriers (Figure 1a-v). Two CpGs were within the proximal enhancer (cg08915438, cg17030679), one within the transcription start site (cg20813374) and the remaining three were within the undefined regions of the *FKBP5* locus (cg6087101, cg16052510, cg23713875; Figure 1a-v). No associations with genotype were replicated in cohort 2.

After FDR correction, several CpGs were significantly altered by both ageing and the CT/TT risk allele in cohort 1. In the DLPFC we found that five CpGs showed convergence of both ageing and genotype effects (Figure 1b). Two of these CpGs were within the proximal enhancer (cg08915438, cg17030679), one within the transcription start site (cg20813374), and three within an undefined region of the *FKBP5* locus (cg06087101, cg16052510, cg23713875). Replication in the OFC did not identify any conserved CpGs significant for the convergence of both age and genotype effects on DNAm in *FKBP5* (Figure 1b).

With these functionally annotated CpGs, we sought to determine potential consequences of the DNAm landscape across the *FKBP5* locus. To understand what downstream physiological processes may be modulated by differential DNAm in *FKBP5*, we wanted to discern whether any transcription factors had binding regions overlap with CpGs within the *FKBP5* locus. Using the ReMap2022 ChIP-seq database and limiting to studies using human cortical tissue [14] [GSE129039 [46] and GSE116825 [27]], we identified three transcription factor binding sites that fell within the *FKBP5* locus: CTCF (Figure 1a-vi), OLIG2 (Figure 1a-vii) and SIN3A (Figure 1a-viii). These binding sites fell predominantly within the functionally annotated regions of *FKBP5*, with all three transcription factors binding across both the proximal enhancer and transcription start site. Two of the four replicated ageing associated CpGs fell within known transcription factor binding sites, specifically cg18468088 (distal TAD) in a CTCF binding site and cg14284211 (intron 5) in an OLIG2 binding site (Figure 1a-iv).

### Altered DNAm across *FKBP5* associated with neuronal cell proportion

Our recent work identified that ageing and disease effects on *FKBP5* expression (mRNA and protein) are most evident in superficial neurons of the neocortex [31]. As cell type is considered one of the strongest influences on DNAm variance [16], we sought to understand whether cell type was associated with differential DNAm across the *FKBP5* locus. Without considering diagnosis and after FDR correction, there were considerable effects to DNAm associated with the proportion of pan-neuronal marker positive (NeuN+) cells. Specifically, 22 of the 50 (44%) analysed CpGs within the TAD boundaries of *FKBP5* were significantly associated with NeuN proportions (Figure 1b). As with age and genotype, the proximal enhancer was the primary contributor of significant CpGs, with seven CpGs falling within the bounds of this region (cg23416081, cg06937024, cg17030679, cg19226017, cg08915438, cg07944278, cg13719443). The NeuN+ cell proportion was generally negatively associated with DNAm in the proximal enhancer (5/7 hits). Of the 16-remaining significant CpGs, 11 fell across the five other functional regions of the *FKBP5* locus: one within the distal TAD (cg05593667), one within the downstream conserved region (cg04192760), two within intron 5 (cg07633853, cg14284211), four within the transcription start site (cg16012111, cg01294490, cg20813374, cg00130530) and two within the proximal TAD (cg10780318, cg01321308). For these regions, an increased proportion of NeuN+ cells associated with greater DNAm. The remaining 5 significant CpGs fell within the undefined regions of *FKBP5* (cg03566752, cg06087101, cg07061368, cg09009770, cg15929276). In the replication cohort, nine CpGs were associated with neuronal proportion, with seven of these replicating findings from the DLPFC.

### CpG specific changes to DNAm across *FKBP5* show case-control differences

We next stratified our analysis by diagnostic status to identify how DNAm levels are changed across disorders in cohort 1 (schizophrenia/ major depression/ bipolar disorder *vs* control). Of the three diagnostic groups, altered DNAm across *FKBP5* was associated most strongly with schizophrenia which had a total of nine significantly associated CpGs (Figure 2b). Six fell within the bounds of a functionally annotated region (Figure 2a-iii), including the proximal enhancer (cg08915438, cg26868354), proximal TAD (cg01321308, cg10780318), transcription start site (cg16012111) and downstream conserved region (cg041992760). Five of these CpGs show decreased levels of DNAm compared to controls, two of which fell within the proximal enhancer (Figure 2a-iii). Major depression shared one significant CpG with schizophrenia (cg26868354, proximal enhancer; Figure 2a-iv, 2b), as well as an additional two significant CpGs, one within the downstream conserved region (cg20719122) and one within an undefined region (cg07194798; Figure 2a-iv). Both significant CpGs within the bounds of a functionally annotated region were hypomethylated in major depression compared to controls (Figure 2a-iv). Bipolar disorder showed no CpG specific association with altered DNAm (Figure 2a-v, 2b).

**Figure 2:**
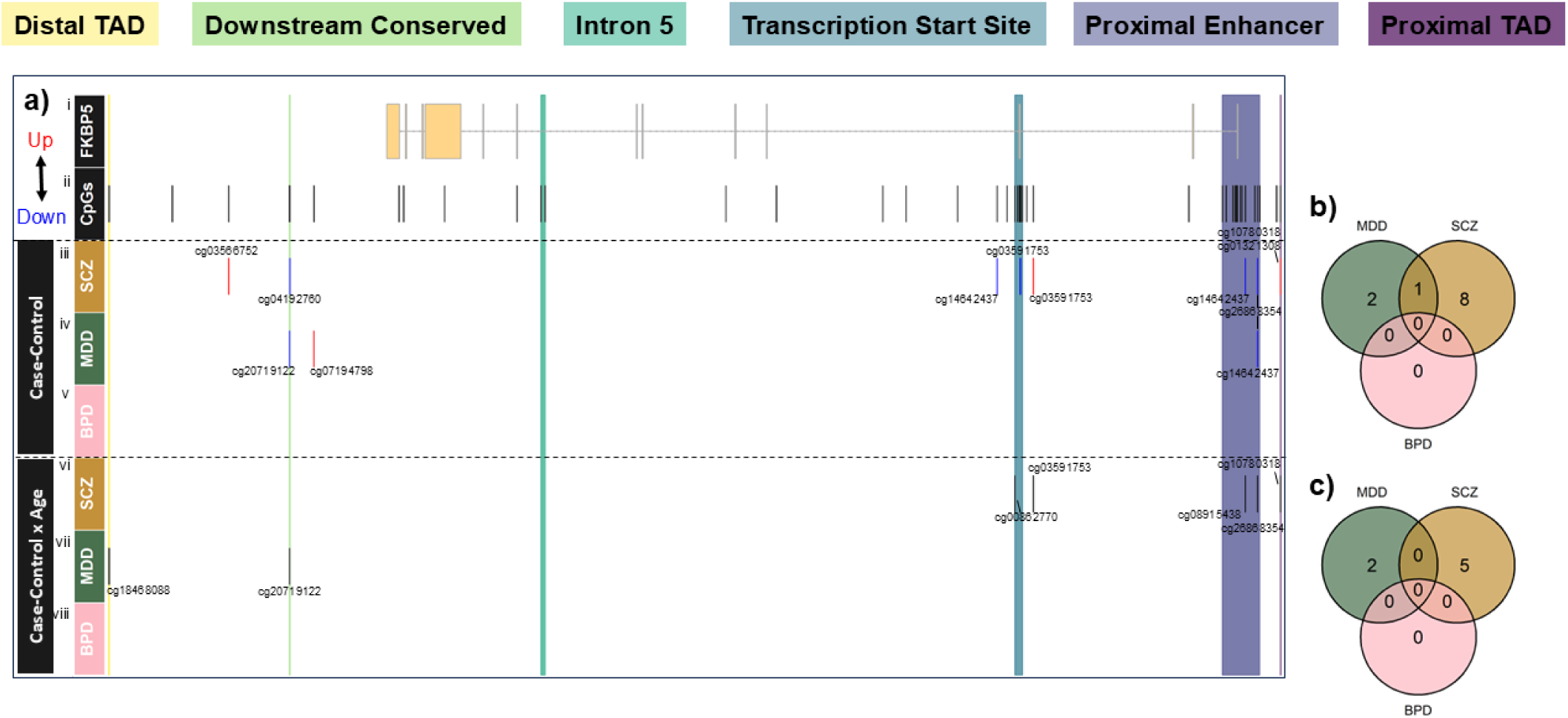
Functionally annotated CpGs in *FKBP5* locus demonstrate diagnosis and ageing trajectory differences, compared to healthy controls. (a) CpGs captured by 450K array, 50kb up/down-stream of the (i) *FKBP5* gene, chromosome 6, were considered for analysis (captured CpGs shown in track). (iii-v) Significantly different CpGs in cases relative to healthy controls in iii) schizophrenia, iv) major depression, and v) bipolar disorder (P_FDR_<0.05). Linear regression correcting for age, sex, genotype (rs1360780 CC vs CT/TT), brain pH, postmortem interval (PMI), estimated proportion of neuronal cells (NeuN+), the first four principal components estimated from the background probes (negative control), and the principal components of genotype (ancestry) that affected the main model was used. Red indicates increase in cases, and blue indicates decrease. (vi-viii) Significantly different ageing trajectories (limited from 14 to 80 years of age) in cases relative to healthy controls in viii) schizophrenia, ix) bipolar disorder, and x) major depression (P_FDR_<0.05). *Sm.ancova* using regression values correcting for covariates as above excluding age. Regions highlighted indicate manually annotated functional domains associated with *FKBP5*. **(b)** Venn diagram of significantly different CpGs between cases and controls. **(c)** Venn diagram of CpGs with significantly different age trajectories between cases and controls. Abbreviations: BPD – bipolar disorder, CpG – cytosine-phosphate-guanine, MDD – major depression, SCZ – schizophrenia, TAD – topographically-associated domain.

### Altered DNAm in schizophrenia is associated with altered age trajectory

Given we identified age to be strongly associated with altered *FKBP5* DNAm, we next compared nonparametric regression curves of age to the residuals of the model to identify any diverging DNAm between cases and controls across ageing (cohort 1). This provided further support that the effect of age on altered DNAm associated most strongly with schizophrenia, particularly at the proximal end of *FKBP5* (Figure 2a-vi). Of the five significant ageing CpGs identified in schizophrenia case x age analysis (Figure 2c), two were from the proximal enhancer (cg08915438, cg26868354), one from the proximal TAD (cg10780318), one from the transcription start site (cg00862770) and one from a non-functionally annotated region (cg03591753; Figure 2a-vi). Two of these CpGs fell within the bounds of a CTCF binding site (cg26868354, proximal enhancer; cg10780318, proximal TAD; Figure 1a-vi), and cg26868354 (proximal enhancer) fell within a SIN3A binding site (Figure 1a-viii). Unlike in schizophrenia, significant CpGs in major depression fell within the distal regions of *FKBP5* (Figure 2a-vii, 2b). Specifically, one CpG within the downstream conserved region (cg20719122) and one within the distal TAD (cg18468088; Figure 2a-vii) which also fell within the bounds of a CTCF binding site (Figure 1a-vi). Schizophrenia and major depression shared no significant CpGs (Figure 2c). Finally, bipolar disorder had no CpGs which showed significant effects of both age and diagnostic status, and there were no CpGs that were significant across all three diagnostic groups (Figure 2c).

As part of this analysis, we also investigated the potential convergent effects of diagnostic status x genotype, in addition to genotype x age x diagnostic status on DNAm levels across *FKBP5*. In this analysis, we compared within-group effects of rs1360780 (CC vs CT/TT), either using the regression model (genotype) or the nonparametric (genotype x age) test. We found limited genotype effects (Supplementary Figure 2). For diagnosis x genotype, the only significant CpG was in bipolar disorder within the transcription start site (cg20813374; Supplementary Figure 2a). For diagnosis x genotype x age, the schizophrenia group had two significantly altered CpGs (Supplementary Figure 2b). One CpG was unique to the schizophrenia group (cg16052510, undefined region), while the other was common between the schizophrenia and control groups (cg23713875, undefined region; Supplementary Figure 2b). Taken together, this evidence provides further support that DNAm changes with ageing and genotype are exacerbated in cases of schizophrenia.

### DNAm across the proximal and distal TADs of *FKBP5* show diagnosis-specific differences

Given DNAm inversely correlates with CTCF occupancy [48], and that the TAD domains are important CTCF binding sites, we wanted to examine case-control differences in DNAm in these regions. Compared to controls and cases of major depression, cases of schizophrenia showed significantly increased DNAm in the proximal TAD (Figure 3c). Conversely, cases of bipolar disorder showed significantly decreased DNAm in the proximal TAD compared to cases of major depression (Figure 3c). In the distal TAD, both major depression and schizophrenia showed significantly increased DNAm compared to controls (Figure 3d). Cases of bipolar disorder also had significantly increased DNAm compared to cases with major depression (Figure 3d).

**Figure 3:**
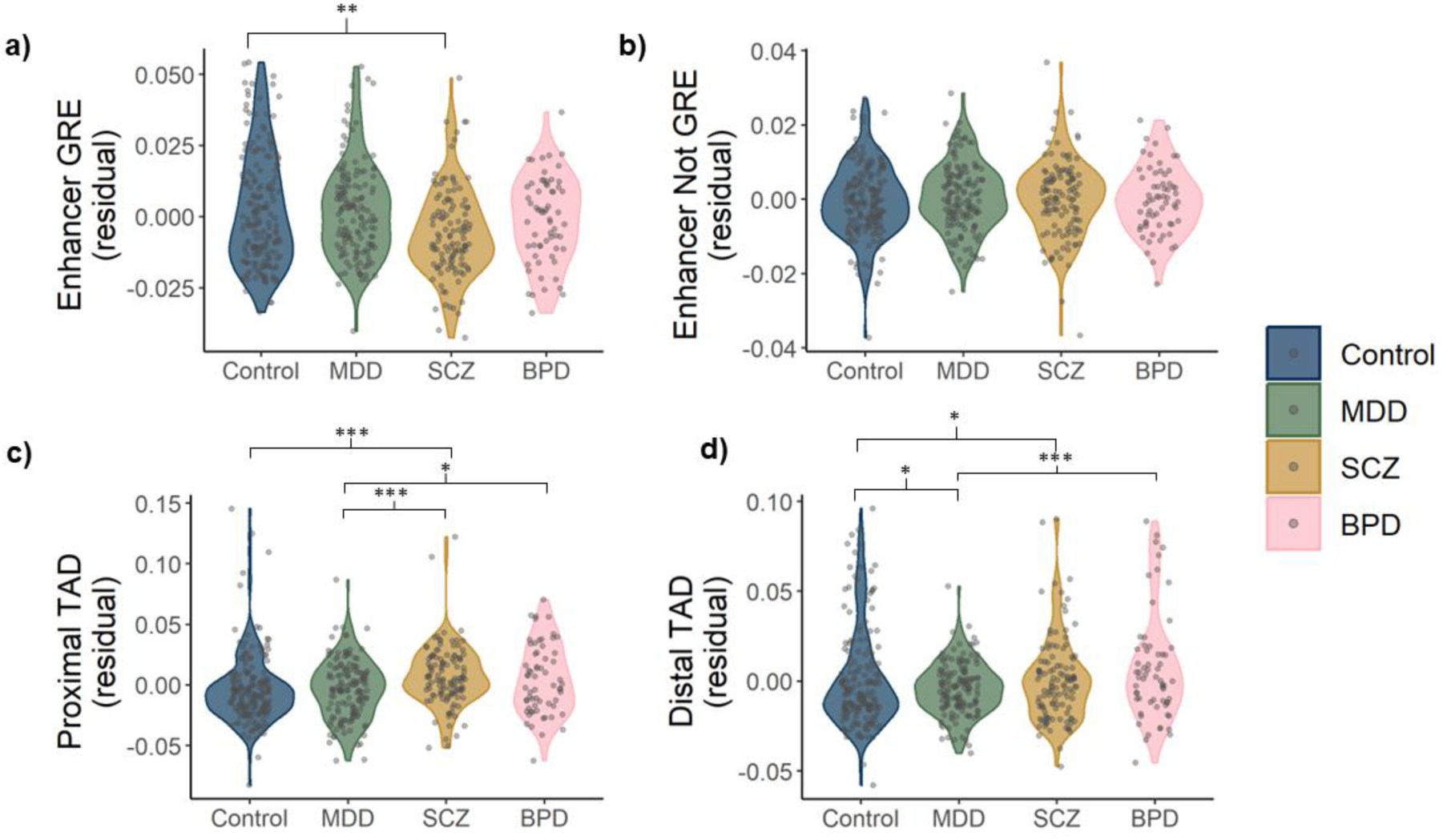
The DNAm of *FBKP5* proximal enhancer and TADs demonstrate robust diagnosis-specific effects on CpGs contained within glucocorticoid response elements. **(a/b)** Case-control regression plots of average CpG methylation proportion in either a) GRE-associated CpGs (n=9) or b) CpGs not associated with GREs (n=5). **(c)** Case-control regression plots of average CpG methylation proportion in the proximal TAD boundary (n=2). **(d)** Case-control regression plots of average CpG methylation proportion in the distal TAD boundary (n=3). Individual residual values from the model are plotted against bulk *FKBP5* gene expression. Significance indicates nominally significant effect associated with diagnosis in the model. *P<0.05, **P<0.01, ***P<0.001. Abbreviations: BPD – bipolar disorder, GRE – glucocorticoid response element, MDD – major depression, SCZ – schizophrenia, TAD – topographically-associated domain.

### Decreased DNAm at key CpGs within proximal enhancer GREs are strongly associated with decreased *FKBP5* gene expression in schizophrenia

Given we consistently observed significant convergent effects of age, genotype and diagnostic status within the proximal enhancer region of *FKBP5*, we next chose to focus on this region to analyse whether CpGs with altered DNAm associate with predicted GREs. Decreased DNAm at key GREs within *FKBP5* is a proposed mechanism by which exposure to early life adversity associates with higher *FKBP5* mRNA induction [22, 23]. When stratified by diagnostic status, we found CpGs within GREs were significantly hypomethylated in schizophrenia compared to controls (Figure 3a). This was not observed for neither major depression nor bipolar disorder, nor was there any significant difference between diagnostic groups and controls for CpGs not within GREs (Figure 3b). This evidence suggests that decreased methylation at GREs within the proximal enhancer bares specific relevance to schizophrenia.

Finally, to determine the influence of methylation patterns on downstream gene function, we examined the relationship between DNAm and *FKBP5* gene expression in cohort 1. Only one CpG (cg00052684) showed significant association when considering gene expression in the main effects model (Figure 4). Further, cg00052684 is located within the proximal enhancer for which decreased methylation was associated with significantly increased *FKBP5* mRNA expression. This trend was reflected across three other CpGs within the proximal enhancer (cg23416081, cg25114611, cg19226017; Figure 4). Interestingly, three of the four CpGs were GRE-linked, notably cg00052684 (Figure 4). This supports our previous findings of increased *FKBP5* induction in BA9, 11 and 24 of older individuals with schizophrenia [31] and suggests the proximal enhancer may be the functional region of *FKBP5* mediating altered expression in response to stress-signalling.

**Figure 4:**
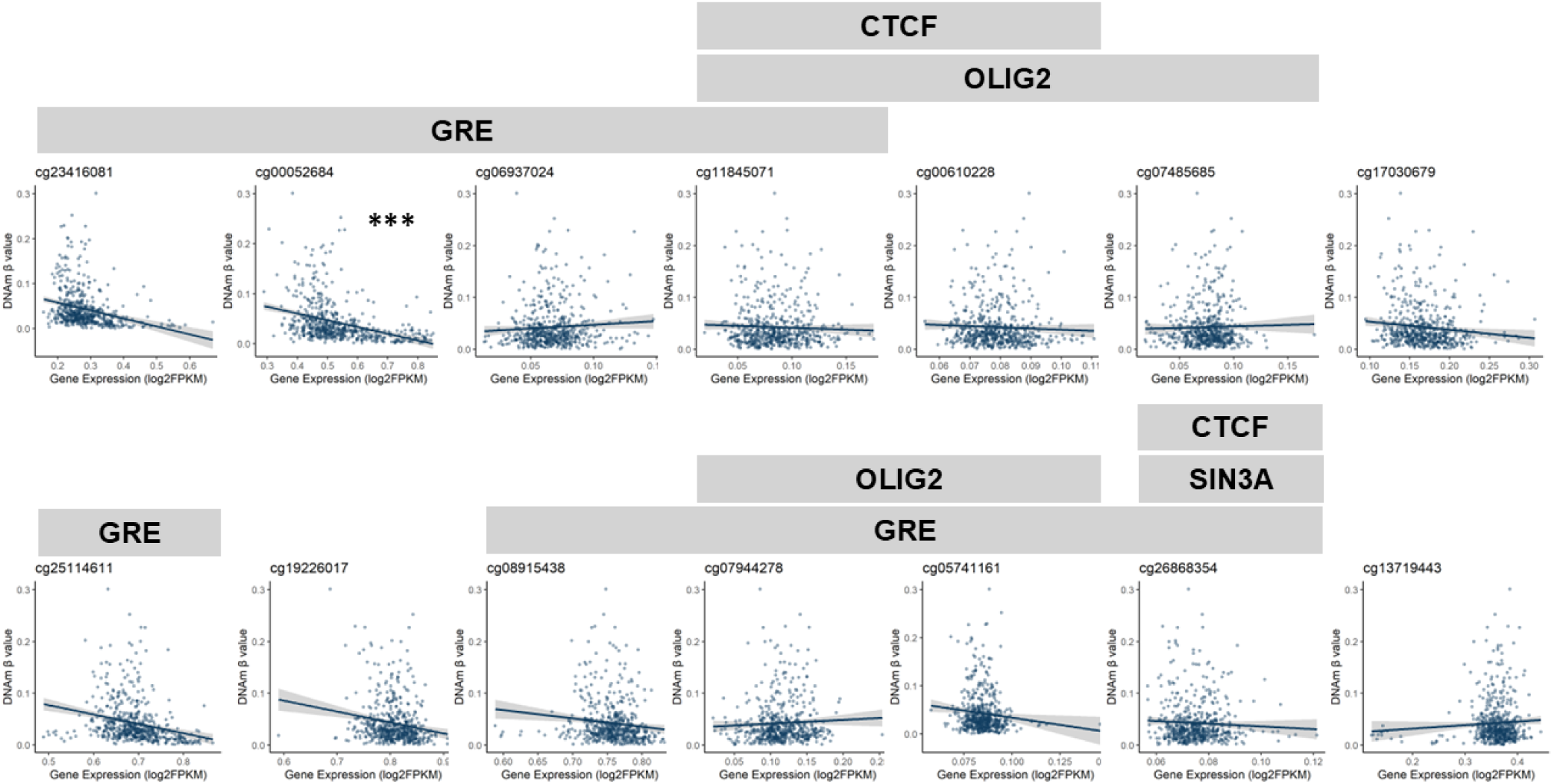
DNAm in glucocorticoid response elements of the proximal enhancer associate with gene expression. Regression plots of *FKBP5* locus CpG B values ∼ log gene expression, correcting for age, sex, postmortem interval, RNA integrity number, pH, genotype, NeuN+ cell proportion, the first four principal components estimated from the background probes (negative control), and the principal components of genotype (ancestry) that affected the main model. Individual residual values from the model are plotted against bulk *FKBP5* gene expression. Grey area indicates 95% confidence interval of model. Outliers > 3 standard deviations were omitted from graphing. Transcription factor binding sites are indicated above CpGs within their binding sites, based on ChIP seq data from cortical datasets ((GSE129039 [46] and GSE116825 [27]). Significance indicates FDR corrected value, *** P_FDR_<0.001. Abbreviations: CTCF – CCCTC-Binding Factor, GRE – glucocorticoid response element, SIN3A – SIN3 Transcription Regulator Family Member A.

## DISCUSSION

We identified altered cortical DNAm at functionally significant regions of *FKBP5* in rs1360780 risk allele carriers and older individuals with schizophrenia, particularly in the TAD boundaries and proximal enhancer of *FKBP5*. We also identified that lower DNAm at GRE-linked CpGs of the *FKBP5* proximal enhancer were associated with increased *FKBP5* mRNA expression. This highlights that modified DNAm in the *FKBP5* locus, particularly in the proximal enhancer and TAD boundaries, may be an important interface for gene-by-environment interactions in the human brain, [30] and supports that gene-by-environment interactions modulate exacerbated trajectories of *FKBP5* expression over the life course in individuals diagnosed with psychiatric disorders [22, 23, 30, 31].

The expression of *FKBP5* increases with ageing [3, 30, 31, 37, 44, 47, 54] and associates with vulnerability for psychiatric disorders, particularly following exposure to early life adversity [6, 30]. When we stratified by diagnosis, schizophrenia disproportionately demonstrated altered DNAm and ageing effects in the DLPFC, particularly in the *FKBP5* proximal enhancer. The proximal enhancer is an important site of tissue-specific transcription factor binding and a key coordinator of gene expression [34]. For *FKBP5* specifically, the proximal enhancer contains a series of stress-responsive GREs whose activation induces transcription [48]. In our study, the GRE-linked CpG cg00052684 in the proximal enhancer was hypomethylated in individuals with schizophrenia in the human cortex, reflecting studies from peripheral systems [22, 23]. There are other CpG-linked GREs across *FKBP5*, including in intron 5 which are altered in peripheral studies, particularly of stress [51]. While we saw hypomethylation with ageing, there were no diagnosis effects. In addition, we identified the proximal enhancer as the primary contributor of age- and genotype-mediated changes to *FKBP5* DNAm. Given the extensive cortical association between increased *FKBP5* mRNA expression and age exacerbated trans-diagnostically in psychopathologies [3, 5, 7, 30, 31, 38, 40, 49], we hypothesise that hypomethylation of GRE-linked CpGs within the proximal enhancer of *FKBP5* may be an upstream point of regulation for *FKBP5* expression in the human cortex. Further, it may represent an epigenetic mechanism by which exposure to early-life adversity can precipitate a trajectory towards psychiatric illness later in life [30, 52].

We also identified that CTCF-linked CpGs in *FKBP5* were hypermethylated in the proximal TAD in schizophrenia and the distal TAD in major depression. CTCF binding is an important coordinator of mammalian 3D genome architecture, regulating gene expression through facilitating interactions of regulatory elements [21, 33, 42]. Increased DNAm at CTCF binding sites can decrease CTCF binding and disrupt chromatin interactions to the point of dysregulated gene expression [9]. Previous studies have shown that binding occupancy of the transcription factor CTCF is inversely correlated to levels of DNAm in *FKBP5* in the periphery [48]. While our findings are congruent with recent work from human blood monocytes showing regulation of *FKBP5* expression by CTCF binding in the TAD boundaries [45], studies in human peripheral blood have also shown that DNAm in these regions are resistant to glucocorticoid signalling [48]. This suggests that in some psychiatric disorders, particularly schizophrenia, increased *FKBP5* expression could be mediated by both hypomethylation at GRE-linked CpGs in the proximal enhancer, as well as altered CTCF binding within the TADs distinctly in the cortex.

Although the size of our postmortem cohort facilitated in-depth analysis of DNAm and *FKBP5* gene expression in the human cortex, there are several methodological considerations. Our previous work showed strong cell-type specific changes to *FKBP5* expression, specifically in schizophrenia [31]. Although DNAm is known to also be cell-type specific [16, 36], here we only rely on estimated proportions of NeuN+ cells per sample, this may be attributable to a dissection artifact, at least in part [39]. Analysis of bulk tissues can also dilute cell-type specific effects, which may be better explored through higher resolution techniques such as single nucleus ATAC-seq. Although significant (FDR-corrected), the effect sizes from this study are small, and thus the biological relevance would need to be further validated. Therefore, a deep understanding of cell-type specific changes to DNAm across *FKBP5* is needed before we can fully explore analysis of *FKBP5* DNAm as a biomarker of stress dysregulation and psychiatric illness [30, 52]. The replication cohort was also notably smaller and from an adjacent cortical region (OFC, BA11) which may reflect the limited number of DNAm effects conserved between cohorts, particularly those associated with diagnosis and rs1360780 genotype.

## CONCLUSIONS

Taken together, our results support that hypomethylation of GRE-linked CpGs within the proximal enhancer of *FKBP5* may have lasting effects on *FKBP5* gene expression in the human cortex. This is particularly relevant to older individuals with schizophrenia, and potentially those who have experienced severe adversity early in life. Further, we hypothesise that diagnosis-specific changes to DNAm in the TAD regions of *FKBP5*, could cause altered CTCF binding, leading to dysregulated *FKBP5* expression and an accelerated trajectory towards psychiatric illness. When considered with previous data, this evidence supports a mechanism of diagnosis-specific *FKBP5* regulation, specifically convergent effects of genotype and age that regulate *FKBP5* at the CpG-specific level. This may have implications for the development of *FKBP5*-targeted therapeutics and further defining a subgroup of patients who will most benefit from such biologically based treatments.

## ABBREVIATIONS

CpGs: cytosine-phosphate-guanine-dinucleotides
CTCF: CCCTC-binding factor
DLPFC: dorsolateral prefrontal cortex
DNAm: DNA methylation
FDR: false discovery rate
FKBP51: FK506-binding protein 51
GR: glucocorticoid receptor
GREs: glucocorticoid response elements
HWE: Hardy-Weinberg equilibrium
NeuN+: pan-neuronal marker positive
OFC: orbitofrontal cortex
PFC: prefrontal cortex
PMI: postmortem interval
qSVA: quality surrogate variable analysis
SNP: single nucleotide polymorphism
TADs: topologically associating domains

## DECLARATIONS

### ETHICS APPROVAL

560 postmortem human brains were acquired by the Lieber Institute for Brain Development Repository via material transfer agreement (Table 1, cohort 1). Brain tissue was donated at the time of autopsy from legal next of kin using audiotaped witnessed informed consent, through the NIMH Human Brain Collection Core under NIMH/NIH Institutional Review Board protocol #90-M-0142, and through the National Institute of Child Health and Human Development Brain and Tissue Bank for Developmental Disorders (http://www.BTBank.org/), under contracts NO1-HD-4-3368 and NO1-HD-4-3383, approved by the Institutional Review Board of the University of Maryland. A replication cohort of 91 subjects was acquired from the NSW Brain Tissue Resource Centre (University of Sydney, Australia; Table 1, cohort 2). The study was approved by the Ludwig Maximilian University Ethics Committee (project 17-085, application 22-0523) and prospective informed consent by the donor, or legal next of kin was obtained for each subject.

### CONSENT FOR PUBLICATION

Not applicable.

### DATA AVAILABILITY

The datasets used and/or analysed during the current study are available from the corresponding author on reasonable request.

### CONFLICT OF INTREST

Dr Binder is a co-inventor of the following patent applications: *FKBP5*: a novel target for antidepressant therapy. European Patent # EP1687443 B1: Polymorphisms in ABCB1 associated with a lack of clinical response to medicaments. United States Patent # 8030033; Means and methods for diagnosing predisposition for treatment emergent suicidal ideation (TESI). European application number: 08016477.5, international application number: PCT/EP2009/061575. The remaining authors declare no competing interests or conflicts of interest.

### AUTHOUR CONTRIBUTIONS

KE and DK performed the analyses, drafted and revised the work. NM and EB conceived, designed and managed the work, performed all analyses and contributed to drafting the manuscript. NY and DC were substantially involved in conceptualising and assisting with the statistical analyses. RT, AJ, JK, DW, TH provided the data from cohort 1, NY and EB contributed the data from cohort 2. NM, EB, AF, JK, DW and TH were involved in the dissection and curation of the brain samples used in this study, as well as managing the experiments and sharing the resulting datasets. All authors were involved in the interpretation of the data and approved the final version of the manuscript.

## ACKNOWLEDGMENTS

We thank the families from the USA and Australia who donated brain tissues to this research. We specifically thank the Lieber and Maltz families for their contributions towards the Lieber Institute brain collection. Tissues received from the New South Wales Brain Tissue Resource Centre at the University of Sydney were supported by the University of Sydney. Research reported in this publication was supported by the National Institute of Alcohol Abuse and Alcoholism of the National Institutes of Health under Award Number NIAAA012725-15. The content is solely the responsibility of the authors and does not represent the official views of the National Institutes of Health.

## REFERENCES

1. Aryee MJ, Jaffe AE, Corrada-Bravo H, Ladd-Acosta C, Feinberg AP, Hansen KD, et al. Minfi: a flexible and comprehensive Bioconductor package for the analysis of Infinium DNA methylation microarrays. Bioinformatics. 2014; 30(10):1363–9.

2. Benjamini Y, Hochberg Y. Controlling the False Discovery Rate: A Practical and Powerful Approach to Multiple Testing. J R Stat Series B (Methodological). 1995; 57(1):289–300.

3. Blair LJ, Nordhues BA, Hill SE, Scaglione KM, O’Leary JC, Fontaine SN, et al. Accelerated neurodegeneration through chaperone-mediated oligomerization of tau. J Clin Invest. 2013; 123(10):4158–69.

4. Bowman AW, Azzalini A. (1997) Applied smoothing techniques for data analysis: the kernel approach with S-Plus illustrations. USA: Oxford University Press.

5. Chen H, Wang N, Zhao X, Ross CA, O’Shea KS, McInnis MG. Gene expression alterations in bipolar disorder postmortem brains. Bipolar Disord. 2013; 15(2):177–87.

6. Comasco E, Gustafsson PA, Sydsjö G, Agnafors S, Aho N, Svedin CG. Psychiatric symptoms in adolescents: FKBP5 genotype-early life adversity interaction effects. Eur Child Adolesc Psychiatry. 2015; 24(12):1473–83.

7. Darby MM, Yolken RH, Sabunciyan S. Consistently altered expression of gene sets in postmortem brains of individuals with major psychiatric disorders. Transl Psychiatry. 2016; 6(9):e890.

8. Ferrari AJ, Santomauro DF, Herrera AMM, Shadid J, Ashbaugh C, Erskine HE, et al. Global, regional, and national burden of 12 mental disorders in 204 countries and territories, 1990–2019: a systematic analysis for the Global Burden of Disease Study 2019. Lancet Psychiatry. 2022; 9(2):137–50.

9. Flavahan WA, Drier Y, Liau BB, Gillespie SM, Venteicher AS, Stemmer-Rachamimov AO, et al. Insulator dysfunction and oncogene activation in IDH mutant gliomas. Nature. 2016; 529(7584):110–4.

10. Fröhlich AS, Gerstner N, Gagliardi M, Ködel M, Yusupov N, Matosin N, et al. Single-nucleus transcriptomic profiling of human orbitofrontal cortex reveals convergent effects of aging and psychiatric disease. Nat Neurosci. 2024; 27(10):2021–32.

11. Gaali S, Kirschner A, Cuboni S, Hartmann J, Kozany C, Balsevich G, et al. Selective inhibitors of the FK506-binding protein 51 by induced fit. Nat Chem Biol. 2015; 11(1):33–7.

12. Gassen NC, Chrousos GP, Binder EB, Zannas AS. Life stress, glucocorticoid signaling, and the aging epigenome: Implications for aging-related diseases. Neurosci Biobehav Rev. 2017; 74:356–65.

13. Halse M, Steinsbekk S, Hammar Å, Wichstrøm L. Longitudinal relations between impaired executive function and symptoms of psychiatric disorders in childhood. J Child Psychol Psychiatry. 2022; 63(12):1574–82.

14. Hammal F, Pierre, Bergon A, Lopez F, Ballester B. ReMap 2022: a database of Human, Mouse, Drosophila and Arabidopsis regulatory regions from an integrative analysis of DNA-binding sequencing experiments. Nucleic Acids Res. 2022; 50(D1):D316–D25.

15. Hartmann J, Wagner KV, Gaali S, Kirschner A, Kozany C, Ruhter G, et al. Pharmacological Inhibition of the Psychiatric Risk Factor FKBP51 Has Anxiolytic Properties. J Neurosci. 2015; 35(24):9007–16.

16. Jaffe AE, Irizarry RA. Accounting for cellular heterogeneity is critical in epigenome-wide association studies. Genome Biol. 2014; 15(2):R31.

17. Jaffe AE, Gao Y, Deep-Soboslay A, Tao R, Hyde TM, Weinberger DR, et al. Mapping DNA methylation across development, genotype and schizophrenia in the human frontal cortex. Nat Neurosci. 2016; 19(1):40–7.

18. Jaffe AE, Tao R, Norris AL, Kealhofer M, Nellore A, Shin JH, et al. qSVA framework for RNA quality correction in differential expression analysis. PNAS. 2017; 114(27):7130–5.

19. Kaul D, Smith CC, Stevens J, Frohlich AS, Binder EB, Mechawar N, et al. Severe childhood and adulthood stress associates with neocortical layer-specific reductions of mature spines in psychiatric disorders. Neurobiol Stress. 2020; 13:e100270.

20. Kendler KS, Myers JM, Keyes CLM. The Relationship Between the Genetic and Environmental Influences on Common Externalizing Psychopathology and Mental Wellbeing. Twin Res Hum Genet. 2011; 14(6):516–23.

21. Kim S, Yu N-K, Kaang B-K. CTCF as a multifunctional protein in genome regulation and gene expression. Exp Mol Med. 2015; 47(6):e166-e.

22. Klengel T, Mehta D, Anacker C, Rex-Haffner M, Pruessner JC, Pariante CM, et al. Allele-specific FKBP5 DNA demethylation mediates gene–childhood trauma interactions. Nat Neurosci. 2013; 16(1):33–41.

23. Klengel T, Binder EB. *FKBP5* Allele-Specific Epigenetic Modification in Gene by Environment Interaction. Neuropsychopharmacol. 2015; 40(1):244–6.

24. Lawrence M, Huber W, Pagès H, Aboyoun P, Carlson M, Gentleman R, et al. Software for Computing and Annotating Genomic Ranges. PLoS Comput Biol. 2013; 9(8):e1003118.

25. Leek JT, Johnson WE, Parker HS, Jaffe AE, Storey JD. The sva package for removing batch effects and other unwanted variation in high-throughput experiments. Bioinformatics. 2012; 28(6):882–3.

26. Liao Y, Smyth GK, Shi W. featureCounts: an efficient general purpose program for assigning sequence reads to genomic features. Bioinformatics. 2013; 30(7):923–30.

27. Lu L, Liu X, Huang W-K, Giusti-Rodríguez P, Cui J, Zhang S, et al. Robust Hi-C maps of enhancer-promoter interactions reveal the function of non-coding genome in neural development and diseases. Mol Cell. 2020; 79(3):521–34.

28. Maksimovic J, Phipson B, Oshlack A. A cross-package Bioconductor workflow for analysing methylation array data. F1000Research. 2017; 5:1281.

29. Matosin N, Cruceanu C, Binder EB. Preclinical and Clinical Evidence of DNA Methylation Changes in Response to Trauma and Chronic Stress. Chronic Stress (Thousand Oaks). 2017; 1:1–20.

30. Matosin N, Halldorsdottir T, Binder EB. Understanding the Molecular Mechanisms Underpinning Gene by Environment Interactions in Psychiatric Disorders: The *FKBP5* Model. Biol Psychiatry 2018; 83:821–30.

31. Matosin N, Arloth J, Czamara D, Edmond KZ, Maitra M, Fröhlich AS, et al. Associations of psychiatric disease and ageing with FKBP5 expression converge on superficial layer neurons of the neocortex. Acta Neuropathol. 2023; 145(4):439–59.

32. Nejati V, Majdi R, Salehinejad MA, Nitsche MA. The role of dorsolateral and ventromedial prefrontal cortex in the processing of emotional dimensions. Sci Rep. 2021; 11(1).

33. Ong C-T, Corces VG. CTCF: an architectural protein bridging genome topology and function. Nat Rev Genet. 2014; 15(4):234–46.

34. Pachano T, Haro E, Rada-Iglesias A. Enhancer-gene specificity in development and disease. Development. 2022; 149(11):dev186536.

35. R Core Team. R: A language and environment for statistical computing. Vienna, Austria: R Foundation for Statistical Computing; 2021 [Available from: https://www.R-project.org/].

36. Rahman MF, McGowan PO. Cell-type-specific epigenetic effects of early life stress on the brain. Transl Psychiatry. 2022; 12(1).

37. Sabbagh JJ, O’Leary JC, 3rd, Blair LJ, Klengel T, Nordhues BA, Fontaine SN, et al. Age-associated epigenetic upregulation of the FKBP5 gene selectively impairs stress resiliency. PloS one. 2014; 9(9):e107241.

38. Sinclair D, Fillman SG, Webster MJ, Weickert CS. Dysregulation of glucocorticoid receptor co-factors FKBP5, BAG1 and PTGES3 in prefrontal cortex in psychotic illness. Sci Rep. 2013; 3(1).

39. Tao R, Davis KN, Li C, Shin JH, Gao Y, Jaffe AE, et al. GAD1 alternative transcripts and DNA methylation in human prefrontal cortex and hippocampus in brain development, schizophrenia. Mol Psychiatry. 2018; 23(6):1496–505.

40. Tatro ET, Everall IP, Masliah E, Hult BJ, Lucero G, Chana G, et al. Differential Expression of Immunophilins FKBP51 and FKBP52 in the Frontal Cortex of HIV-Infected Patients with Major Depressive Disorder. J Neuroimmune Pharmacol. 2009; 4(2):218–26.

41. Teschendorff AE, Marabita F, Lechner M, Bartlett T, Tegner J, Gomez-Cabrero D, et al. A beta-mixture quantile normalization method for correcting probe design bias in Illumina Infinium 450 k DNA methylation data. Bioinformatics. 2013; 29(2):189–96.

42. Thiecke MJ, Wutz G, Muhar M, Tang W, Bevan S, Malysheva V, et al. Cohesin-dependent and-independent mechanisms mediate chromosomal contacts between promoters and enhancers. Cell Rep. 2020; 32(3).

43. Visscher PM, Wray NR, Zhang Q, Sklar P, McCarthy MI, Brown MA, et al. 10 Years of GWAS Discovery: Biology, Function, and Translation. Am J Hum Genet. 2017; 101(1):5–22.

44. Volk N, Julius, Engel M, Anthony, Cattane N, Cattaneo A, et al. Amygdalar MicroRNA-15a Is Essential for Coping with Chronic Stress. Cell Rep. 2016; 17(7):1882–91.

45. Wang C, Manders F, Groh L, Oldenkamp R, Logie C. Corticosteroid-induced chromatin loop dynamics at the *FKBP5* gene. Ann N Y Acad Sci. 2023; 1529(1):109–19.

46. Wang Q, Zhao Z, Xu H, Lu D, Xia J, Zhang W, et al. Computational divergence analysis reveals the existence of regulatory degeneration and supports HDAC1 as a potential drug target for Alzheimer’s disease. bioRxiv. 2023:2023.10. 05.561015.

47. Weickert CS, Webster MJ, Boerrigter D, Sinclair D. *FKBP5* Messenger RNA Increases After Adolescence in Human Dorsolateral Prefrontal Cortex. Biol Psychiatry. 2016; 80(5):29–31.

48. Wiechmann T, Röh S, Sauer S, Czamara D, Arloth J, Ködel M, et al. Identification of dynamic glucocorticoid-induced methylation changes at the FKBP5 locus. Clin Epigenetics. 2019; 11(1):83.

49. Young KA, Thompson PM, Cruz DA, Williamson DE, Selemon LD. BA11 FKBP5 expression levels correlate with dendritic spine density in postmortem PTSD and controls. Neurobiol Stress. 2015; 2:67–72.

50. Young SG, Bowman AW. Non-parametric analysis of covariance Biometrics. 1995; 51:920–31.

51. Yusupov N, Roeh S, Sotillos Elliott L, Chang S, Loganathan S, Urbina-Treviño L, et al. DNA methylation patterns of FKBP5 regulatory regions in brain and blood of humanized mice and humans. Mol Psychiatry. 2024; 29(5):1510–20.

52. Zannas AS, Wiechmann T, Gassen NC, Binder EB. Gene–Stress–Epigenetic Regulation of FKBP5: Clinical and Translational Implications. Neuropsychopharmacol. 2016; 41(1):261–74.

53. Zannas AS, Jia M, Hafner K, Baumert J, Wiechmann T, Pape JC, et al. Epigenetic upregulation of FKBP5 by aging and stress contributes to NF-kappaB-driven inflammation and cardiovascular risk. Proc Natl Acad Sci U S A. 2019; 116(23):11370–9.

54. Zheng D, Sabbagh JJ, Blair LJ, Darling AL, Wen X, Dickey CA. MicroRNA-511 Binds to FKBP5 mRNA, Which Encodes a Chaperone Protein, and Regulates Neuronal Differentiation. JBC. 2016; 291(34):17897–906.

