## Supplementary Materials for "Lower cortical *FKBP5* DNA methylation at key enhancer sites is associated with older age and higher gene expression in schizophrenia"

**Supplementary Table 1: Annotated CpGs based on functional domain of the *FKBP5* locus**

| | Chrom Start | CpG | Annotated Region |
|---|---|---|---|
| | 35490743 | cg05593667 | Distal TAD |
| | 35490787 | cg02837122 | |
| | 35490817 | cg18468088 | |
| | 35523699 | cg20719122 | Downstream Conserved |
| | 35523734 | cg04192760 | |
| | 35569471 | cg07633853 | Intron 5 |
| | 35570223 | cg14284211 | |
| | 35655763 | cg00862770 | Transcription Start Site |
| | 35656242 | cg00140191 | |
| | 35656589 | cg10913456 | |
| | 35656757 | cg16012111 | |
| | 35656847 | cg07843056 | |
| | 35656905 | cg01294490 | |
| | 35657180 | cg20813374 | |
| | 35657202 | cg00130530 | |
| | 35693572 | cg23416081 | Proximal Enhancer |
| | 35694245 | cg00052684 | |
| | 35695489 | cg06937024 | |
| | 35695859 | cg11845071 | |
| | 35695933 | cg00610228 | |
| | 35696060 | cg07485685 | |
| | 35696299 | cg17030679 | |
| | 35696870 | cg25114611 | |
| | 35697184 | cg19226017 | |
| | 35697759 | cg08915438 | |
| | 35699424 | cg07944278 | |
| | 35699498 | cg05741161 | |
| | 35699951 | cg26868354 | |
| | 35700381 | cg13719443 | |
| | 35704148 | cg10780318 | Proximal TAD |
| | 35704223 | cg01321308 | |

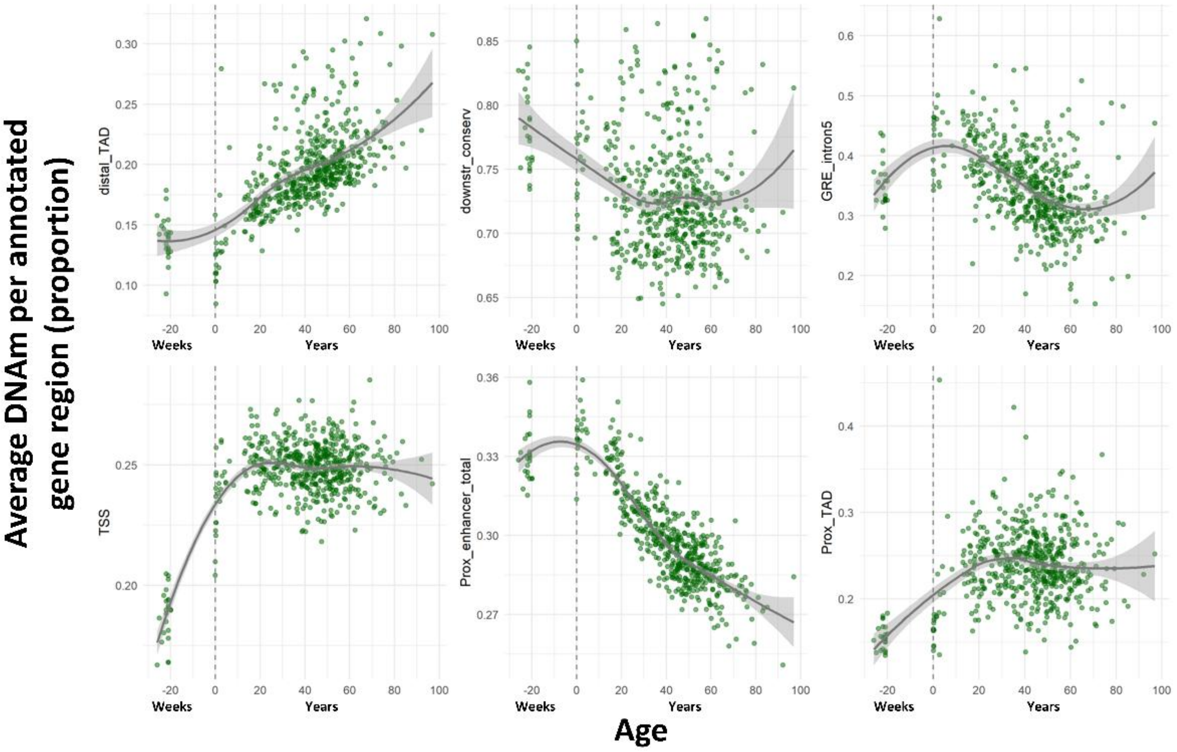


**Supplementary Figure 1: Ageing patterns of DNAm in functionally annotated domains across the healthy lifespan.** Mean DNAm beta value per functionally annotated region per individual across the extended healthy cohort 1 (n=180 + 51). Trendline indicates *loess* fit line with shaded area indicating 95% confidence interval. On x-axis, weeks indicates weeks prenatal, and years indicates years postnatal. Dashed line indicates age=0. Abbreviations: GRE – glucocorticoid response element, TAD – topographically-associated domain, TSS – transcription start site.

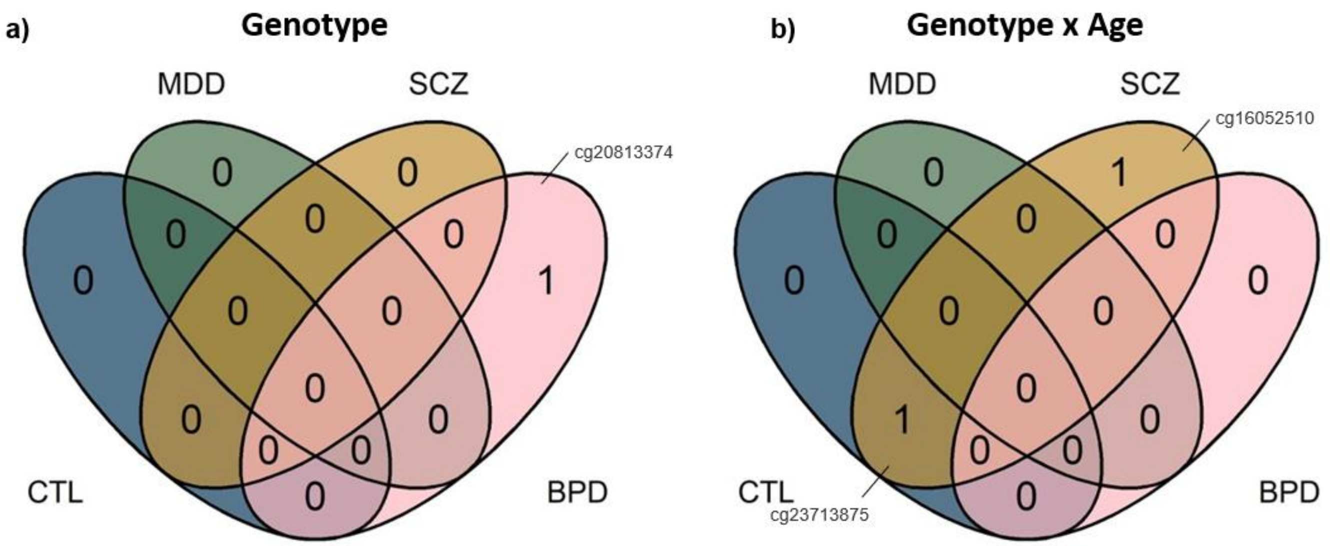


**Supplementary Figure 2: Genotype and genotype x age demonstrate limited diagnosis-specific association. (a)** Venn diagram of CpGs significantly different between CC vs CT/TT genotypes of rs1360780 within each diagnosis. **(b)** Venn diagram of CpGs with significantly different ageing trajectories between CC vs CT/TT rs1360780 genotypes within each diagnosis. Significance indicates FDR corrected P values ($P_{FDR}$<0.05, Benjamini-Hochberg correction). Abbreviations: BPD – bipolar disorder, CTL – control, MDD – major depressive disorder, SCZ – schizophrenia.
